# Stopping a Continuous Movement: A Novel Approach to Investigating Motor Control

**DOI:** 10.1101/2021.04.08.439070

**Authors:** Kelsey E. Schultz, Dominique Denning, Vanessa Hufnagel, Nicole Swann

## Abstract

Flexible, adaptive behavior is critically dependent on inhibitory control. For example, if you suddenly notice you are about to step on a tack and would prefer not to, the ability to halt your ongoing movement is critical. To address limitations in existing approaches for studying your ability to rapidly terminate your movement (“stopping”), we developed a novel stop task. This task requires termination of ongoing motor programs, provides a direct measure of SSRT, and allows for comparison of the same behavior (stopping) in conditions that elicit either prepared or reactive inhibitory control. Here, we present and evaluate our novel Continuous Movement Stop Task (CMST). We examined several versions of the task in a total of 49 participants. Our data reveal that the CMST is effectively able to dissociate stopping behavior between the planned and unplanned conditions. Additionally, within the subset of participants for which we measured speed, we found that participants initiated stopping (with respect to the stop signal) significantly earlier on planned stop compared to unplanned stop trials. Finally, participants took longer to arrive at full motor arrest (i.e. SSRT) following stop initiation on planned than on unplanned stop trials. This novel task design will enable a more precise quantification of stopping behavior and, in conjunction with neuroscientific methods, could provide more rigorous characterization of brain networks underlying stopping.

## INTRODUCTION

Our ability to behave in a flexible, adaptive manner is critically dependent on inhibitory control. For example, imagine walking barefoot to your refrigerator when you suddenly spot a scorpion on the floor in front of you. As the goal changes from getting food to not getting stung, the ability to halt your ongoing movement is vital. Evidence suggests that the inhibitory processes underlying motor termination, similar to that described in this example, are also important for other forms of cognitive and behavioral control such as emotion regulation (Lewis et al., 2006), attentional control (Zavala et al., 2017), and impulse control (Schachar & Logan, 1990). Disruption of the inhibitory control system may contribute to myriad neurological and psychiatric disorders such as Parkinson’s disease, obsessive-compulsive disorder, schizophrenia, attention-deficit hyperactivity disorder, and substance use disorder (Lipszyc & Schachar, 2010). Given the broad relevance of inhibitory control in degenerative and psychiatric disorders, a comprehensive understanding of this system is essential for development of treatments, interventions, and diagnostic tools.

Much of what we understand about inhibitory processes has been established using standard motor inhibition tasks, such as the stop signal task (SST; Logan & Cowan, 1984). The SST is designed to measure efficiency of response-inhibition by estimating the amount of time required to inhibit an initiated response. Participants are asked to respond as quickly as possible to a ‘go’ signal (typically with a button press) and to inhibit their response when the ‘go’ signal is followed by a ‘stop’ signal. The metric of interest, known as the stop signal reaction time (SSRT), is then derived from go-signal reaction time (RT), stop signal delay, and probability of correct responding (i.e. successfully stopping when prompted). SSRT can be quantified in this way using simple mathematical models, such as the independent race model, which assumes that going and stopping are two independent processes and behavioral output (or lack thereof) is determined by which process is completed first (Logan & Cowan, 1984).

### Measurement of SSRT on Individual Trials

Though the SST has been an important and effective tool for inhibitory control research, there are limitations in its design. One major limiting factor is its inability to unambiguously measure stopping behavior on individual trials. That is, because SSRT is necessarily estimated from several to many trials (see Verbruggen et al., 2019 for best practices), the SST cannot capture variability in the duration of the stopping process across individual trials. Recently a technique was introduced that provides an estimation of SSRT on individual trials using subthreshold electromyography activity as a marker of action cancellation (Jana et al., 2020; Raud & Huster, 2017). However, because this technique requires partial responses, it is only effective for about half of all successful stop trials. Additionally, the SST presents challenges in identifying which stop trials are truly successful versus unsuccessful. For example, on any failed stop trial, it is unclear whether failure to stop is indicative of an initiated stop process that was slower than the go process or if the stop process was never initiated (‘trigger failure’) because in either case the motoric output is identical (failure to stop; Matzke et al., 2017; Verbruggen et al, 2019). Similarly, an apparent successful stop is indistinguishable from a failure to engage with the task at all on that trial. That is, it is not clear whether participants successfully inhibited an already initiated action or if they simply failed to initiate any action at all. This ambiguity complicates interpretation of task performance and, when used in conjunction with neuroscientific techniques, identification of neurophysiological correlates of inhibitory processes. This may be especially problematic for patient or pediatric populations for whom behavioral performance is likely to be more variable. Furthermore, because identifying neurophysiological substrates of stopping with the SST typically requires comparison of successful stop trials with unsuccessful stop trials and/or go trials (i.e. comparing stillness to movement), dissociating neurophysiological correlates of motor termination, prepared inhibitory processes, and sensory feedback from movement (or lack thereof) is challenging, if not impossible.

### Prepared vs. Reactive Stopping

Additionally, the SST is not well suited for interrogating processes underlying prepared motor termination. This is an important line of inquiry, as it has been suggested that prepared inhibitory processes and reactive inhibitory processes (i.e. stopping in reaction to an unexpected stimulus) may recruit different neural mechanisms. That is, reactive inhibition is thought to recruit the monosynaptic ‘hyperdirect’ cortico-basal ganglia pathway, while prepared inhibition may recruit the multisynaptic ‘indirect’ pathway (Aron et al., 2016; Jahfari et al., 2012; Lofredi et al., 2020). The SST can be modified to better suit this line of inquiry by giving participants a cue about the probability that they will be given a stop signal on the next trial (Greenhouse et al., 2012; Muralidharan et al., 2019; Swann et al., 2013). The foreknowledge that a stop signal may occur allows subjects to proactively engage inhibitory mechanisms that will be employed to countermand triggering of the response tendency. However, this approach still suffers from some of the limitations described above, including that one cannot fully disentangle planning to stop an initiated movement and not planning to initiate a movement. Studies employing this technique have found that prepared inhibition is associated with both shorter SSRT (faster stop) and longer RT (slower response), suggesting involvement of a preparatory step prior to the response onset (Aron et al, 2011). The behavioral manifestation of preparing to stop an ongoing movement is as yet unclear.

### Stopping a Continuous Movement

Lastly, the ballistic nature of the movement being suppressed (often a button press) limits generalizability of task performance for stopping an ongoing movement. Although executing a continuous movement recruits different regions than a discrete movement does (Habas & Cabanis, 2008), it is often assumed that both movements recruit the same inhibitory network to implement stopping. This is based on previous work which shows that stopping a variety of movements including button presses, foot movements, speech, and eye movements all seem to engage the same brain networks (Cai et al., 2012; Coe & Munoz, 2017; Greenhouse et al., 2012). However, these approaches are all built on a framework of discrete movements and generalizability to continuous movement that has not yet been established empirically.

Here, we test and describe the performance characteristics of our novel version of the stop signal task that requires termination of a continuous movement (i.e. a circular rotation with a computer mouse) in response to a stop signal under conditions that elicit either proactive or reactive stopping processes. Because participants are stopping an ongoing action on every trial, our continuous movement stop task (CMST) provides a direct measure of individual trial SSRT, rather than an estimate of SSRT over many trials. Additionally, in contrast to methods that require electromyography to identify the time of motor inhibition on individual trials (Jana et al., 2020; Raud & Huster, 2017), SSRT can be measured from behavior alone. The precision offered by this novel design provides a straightforward interpretation of behavioral performance as well as the ability to directly compare prepared to reactive motor termination and investigate stopping processes underlying termination of a continuous movement. Furthermore, it provides an opportunity to probe motor termination using neuroscientific methods (such as electrophysiology and functional imaging) with improved ability to disambiguate motoric and cognitive contributions to stopping, allowing for a clearer examination of the underlying mechanism.

## METHODS

We designed and evaluated a novel continuous movement stop task (CMST) that includes two trial types: unplanned stop trials that probe predominantly reactive stopping processes and planned stop trials that probe predominantly prepared stopping processes. Four versions of this task were performed. The first three versions varied in the percentage of unplanned stop trials relative to planned stop trials. The fourth version included a guide (paper circle attached to the desk) that encouraged limited movement magnitude. This addition allowed us to capture mouse coordinates more reliably, which permitted more accurate speed calculations (a limitation to versions 1-3). We evaluated reaction times to the go and stop signals for all versions and evaluated speed for the fourth version.

### Participants

We collected data from 49 right-handed university students (30 females, 19 males) between the ages of 18 and 32 (*M* _*age*_ = 20.5, *SD* = 0.55). Each participant completed one of four task versions (table 1). Participants were recruited through the university’s online human subjects pool using SONA-systems and through flyers posted throughout campus. All participants signed a consent form approved by the university ethics board.

**Table 1.** Version specific information. Each version was performed by a different group of participants (total N = 49).

| Version | Unplanned Stop Trials | Movement | N | Mean Age | Sex Breakdown |
| --- | --- | --- | --- | --- | --- |
| 1 | 10% (24 total) | No Guide | 13 | 19.8 | 6 females, 7 males |
| 2 | 30% (72 total) | No Guide | 14 | 21.1 | 10 females, 4 males |
| 3 | 50% (120 total) | No Guide | 14 | 20.5 | 7 females, 7 males |
| 4 | 30% (72 total) | Includes Guide | 8 | 20.75 | 7 females, 1 male |

### Task

The CMST (fig. 1) required participants to move a computer mouse in a continuous circular motion (at a rate of about one complete circle per second) while monitoring a countdown that appeared on the screen in front of them. The cursor was not visible during the countdown. The countdown started at any value from 6-3 (with equal probability across both trial types) and counted down to 1 at a regular rate of one number per second. Participants were asked to begin moving as quickly as possible after the go signal was presented and to stop moving as quickly as possible after the stop signal. On some trials, the countdown reached “1” before the stop signal appeared. On the remaining subset of trials, the stop signal appeared at a pseudorandom time-point before the countdown reached “1” (unplanned-stop). Approximate trial length varied between 7 and ∼2.5 seconds and was defined for all trials at the beginning of the task such that both trial types had an equal number of trials of comparable lengths. Stop signal arrival time points for unplanned stop trials were generated using Matlab’s random number generator (*rand(N)*) plus approximate intended trial length. For example, on an unplanned stop trial intended to be approximately 5 seconds long, the stop signal presentation time would be 5 plus a random number between 0 and 1. Arrival of the stop signal during these trials was therefore not predictable. The task was programmed using the Matlab Psychophysics toolbox (Brainard, 1997).

**Figure 1.**
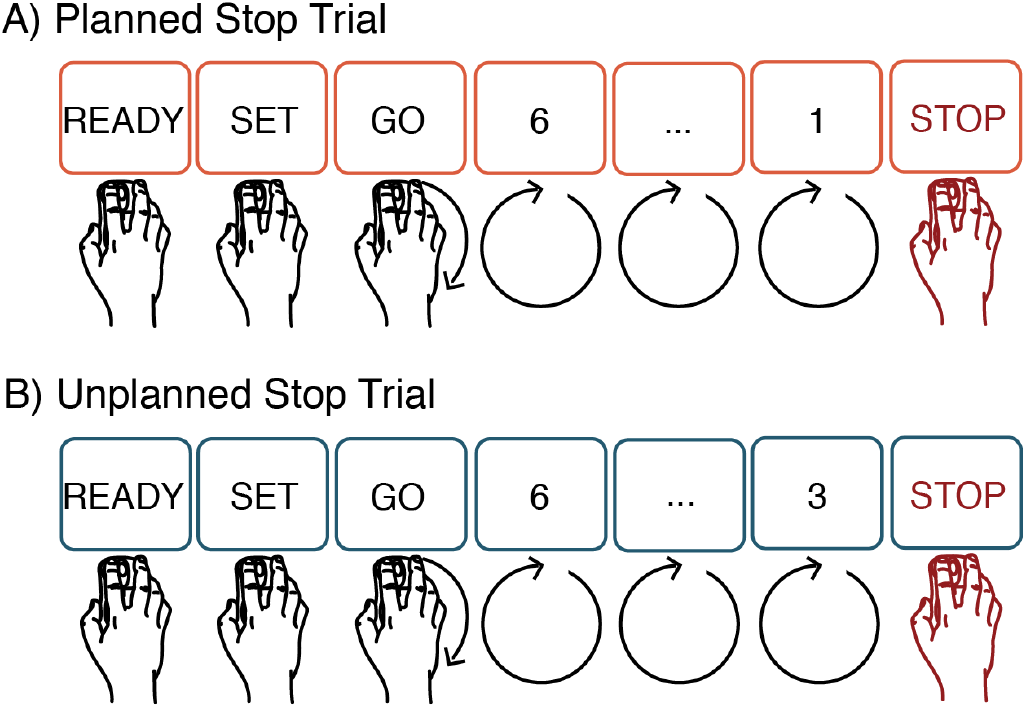
Depiction of planned and unplanned stop trials in CMST.

### Task training

Participants were required to correctly complete at least 5 practice trials before moving on to the rest of the experiment. If participants had not met a set of criteria after completing 20 trials, they did not move on to the rest of the experiment. At the end of each practice trial, participants were given feedback regarding their behavioral performance. There were 6 possible error types: early start (RT< 0ms), late start (RT>1000ms), early stop (SSRT<0ms), late stop (SSRT>1000ms), too slow (speed of movement<10 pixels/ms) and too fast (speed of movement>60 pixels/ms). If behavior across all metrics fell within the window of acceptable behavior, they were told their performance was “perfect”. If behavioral measures fell outside this window, they were prompted to correct it. For example, if a participant waited too long to respond to the go-cue (RT>1000ms), they were reminded to respond as quickly as possible when presented with the go-cue. Following successful completion of the practice trials, participants were permitted to begin the rest of the experiment. Only one participant was unable to successfully complete 5 of the 20 practice trials and therefore did not advance to the rest of the experiment and was not included in the total participant number.

The task included 12 blocks consisting of 20 trials (240 total trials). To promote consistent performance throughout the task and across participants, feedback similar to that given during the practice round was given at the end of each block. Participants were informed of the number of times they made each of the 6 error types. For version 4, block-wise feedback was associated with a score and a color based on performance across that block along with the number of times each error type was made. For every block, a total of 120 errors could be made (6 error types across 20 trials). Scores reflected percentage of errors made out of total possible errors. Any score above 75% was reported in green and a score of 75 or below was reported in red.

To explore how behavior varied with percentage of unplanned stop trials, we ran 3 separate versions of the task (in 3 separate groups of participants; see table 1). Task versions 1-3 contained 10%, 30%, or 50% unplanned stop trials (table 1). In each version, the remaining trials were planned stop trials. A fourth version of the task containing 30% unplanned stop trials was administered to a separate group of participants. In addition to the feedback provided at the end of the block, this version also incorporated a circular paper guide with a diameter of 10 cm secured to the desk participants used. This gave participants a guide for how large their movements should be and helped constrain movement magnitude.

### Data Collection

Participants were seated comfortably in a chair approximately 61cm from a computer screen. A tri-axial accelerometer was attached to their right (dominant) wrist. (Note that here we used accelerometry to verify behavior recorded by mouse coordinates but recording accelerometry is not necessary for the task). All participants used their right hand to control a computer mouse located to the right of the computer screen. Verbal and written instructions were given prior to the start of the task.

### Data Analysis

#### Reaction Times

Reaction times following the go signal (RT) and stop signal (SSRT) were calculated for each participant using mouse location data (fig. 2a). Psychophysics Toolbox reports mouse coordinates as XY location in pixel space, which is defined by the display parameters (1600×800 pixels) and provides a sampling rate of mouse location of 50Hz. RT was defined as the first time point at which the mouse moved at least 1 pixel from its starting location. SSRT was defined as the time point at which change in mouse location did not exceed 3 pixels for at least 200ms. This value was selected with the consideration that task unrelated movements, such as lowering one’s wrist or elbow to rest on the table after stopping, could be recorded as small changes in mouse location. Visual inspection of accelerometry data was used to confirm accuracy of RT and SSRT for all subjects (fig. 2b).

**Figure 2.**
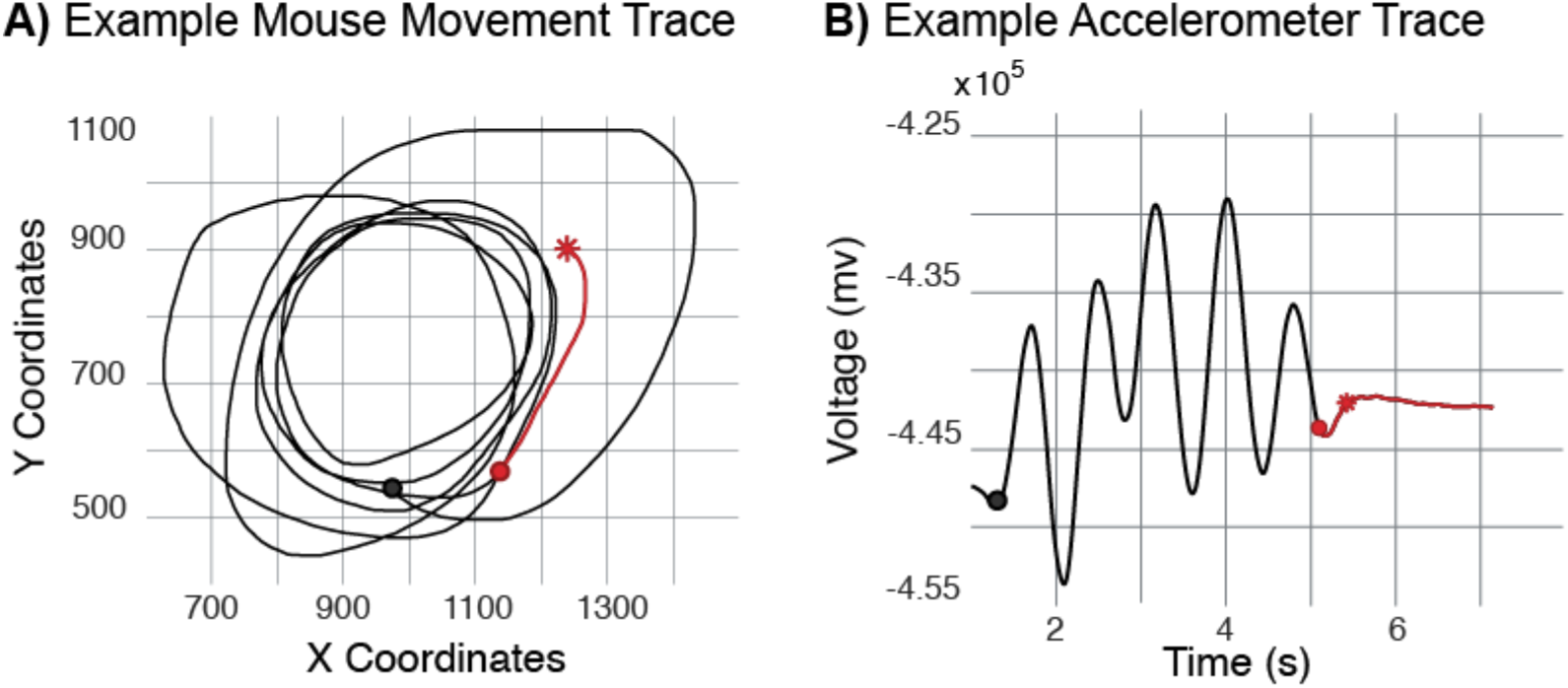
Example Matlab and Accelerometer data from a single trial. Red line indicates time from stop signal presentation (red circle) to movement termination (i.e. SSRT; marked by a red star). A) Changes in computer mouse position recorded with Matlab. Mouse position is continuously recorded throughout a trial as its location in an XY coordinate system defined by monitor size and resolution. Shown here are the changes in XY position between time of go signal presentation (black dot) and SSRT. B) Accelerometer trace for the same trial. Data are shown from go signal presentation (time = 0) to 1.5 seconds after stop signal presentation.

#### Speed

After analysis of the first three versions of the task, we observed that the recorded magnitude of participants’ circular motions frequently exceeded the limits of the coordinate plane used to identify mouse location, resulting in saturation of recorded mouse coordinate data in one dimension (X or Y) for these trials. That is, when movement of the mouse was outside the boundaries defined by the display (1600×900 pixels), it would be recorded as a change in one dimension only (i.e. change in X or Y) rather than a change in both dimensions (i.e. change in X and Y; fig. 3a). On average, the majority (93%) of the trials for versions 1-3 were missing more than 30% of location data and thus were deemed unsuitable for speed calculations. However, this loss of location data did not affect our ability to calculate RT and SSRT since location data was still available for one dimension. Accuracy of RT and SSRT was also confirmed using accelerometry. To better capture behavior related to variations in speed, version 4 included a paper guide to encourage movements that remained within the limits of the coordinate plane and therefore could be recorded more reliably. Speed measures for version 4 were obtained by tracking the change in mouse position in pixel space and were calculated as the rate of change in pixels per millisecond (px/ms). The paper guide was not 100% effective in constraining movement, so participants still exceeded the limits of the coordinate plane on occasion, but much less often than in the previous three versions (fig. 3b). A Bayesian test of equivalence, for task version 4, comparing speed for trials which included out of bounds portions compared to trials which remained entirely in bounds revealed that speed calculations performed on trials with out of bounds data (i.e. missing location data) were not significantly different than speed calculations performed excluding these data points (*BF*_*10*_ = .56) informing our decision to include all trials from version 4 in our speed analyses.

**Figure 3.**
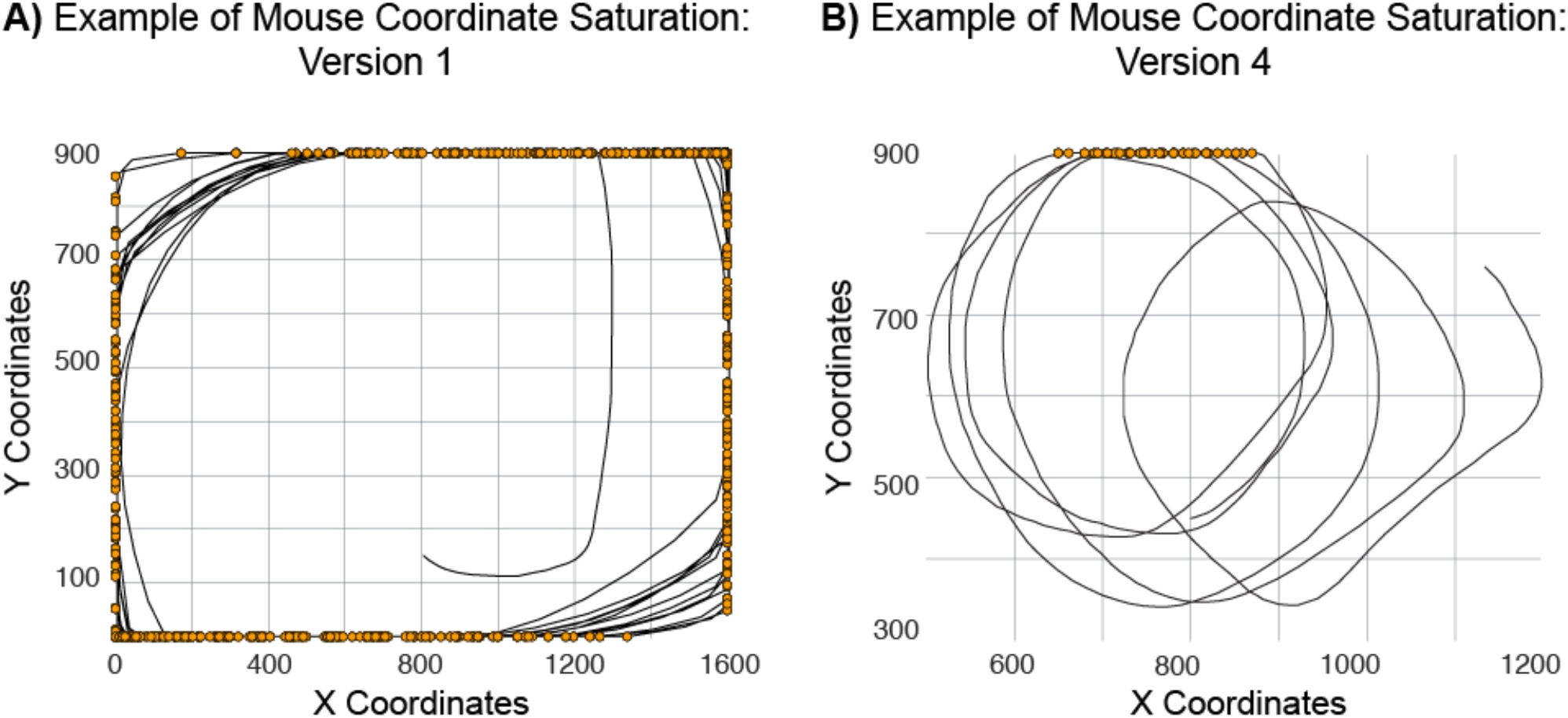
Examples of missing location data. A) Example trial from version 1 in which mouse movement exceeded display boundaries. Yellow dots indicate points at which mouse movement exceeded the lower limits of the Y-axis. Example is representative of trials from versions 1-3. B) Example trial from version 4 in which mouse movement exceeded display boundaries.

Because arrival of the stop signal is predictable on planned stop trials, we expected that participants would slow their movement speed in anticipation of stop signal arrival, but prior to stop signal presentation. In contrast, for unplanned stopping we predicted stopping would be preceded by a slowing period only after stop signal presentation. To identify if participants moved slower in the time immediately before stop signal presentation, we calculated and compared two speed measures: end of trial speed and mid-trial speed. End-of-trial speed was defined as the average speed in the last 500 ms before stop signal presentation. Mid-trial speed was defined as the average speed in a 500 ms time window that ended 100 ms before the start of the end-of-trial window (fig. 4). We then compared the percent change in speed between mid-trial movement and end-of-trial movement. We also calculated the correlation between this percent change in speed and SSRT across trials, since we predicted slowing down in preparation for the stop signal might impact SSRT. Lastly, we compared average speed across the entire trial for each trial type and evaluated the relationship between average trial speed and SSRT.

**Figure 4.**
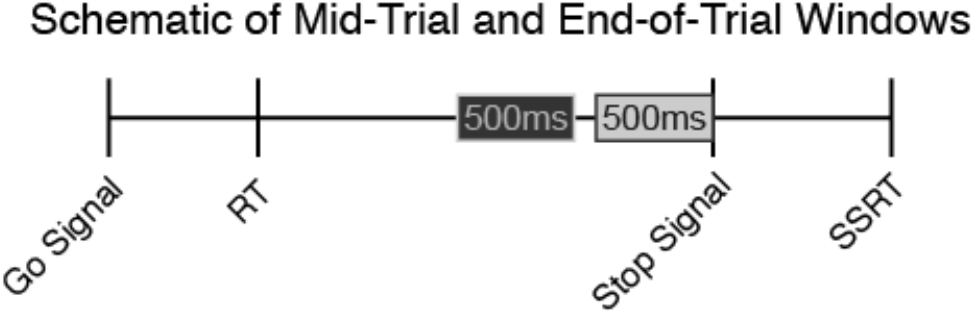
Example definitions of mid-trial and end of trial speed. Dark gray bar indicates mid-trial time window, light gray bar indicates end of trial window.

We also explored the time at which participants began stopping (with respect to stop signal arrival). This measure, henceforth referred to as ‘stop initiation’, was defined as the last time point preceding a consistent deceleration that culminated in a stop (i.e. SSRT; fig. 5). Deceleration was deemed consistent if speed did not subsequently increase by more than 0.5px/ms. We performed 5 analyses/controls using this measure: (1) We assessed differences in stop initiation time between trial types (planned vs unplanned stop trials). (2) We correlated stop initiation with SSRT for both trial types. (3) To evaluate the potential that subjects were moving slower at the time of stop initiation on planned compared to unplanned stop trials we identified instantaneous speed at stop initiation and compared this value between trial types. (4) To examine the possibility that participants were able to stop faster when moving slower we correlated instantaneous speed at stop initiation time with SSRT. Finally, (5) to assess the possibility that the time between stop initiation and full motor arrest (i.e. the amount of time it took participants to stop) was different between planned and unplanned stop trials we took the difference between stop initiation time and SSRT and compared this measure between trial types.

**Figure 5.**
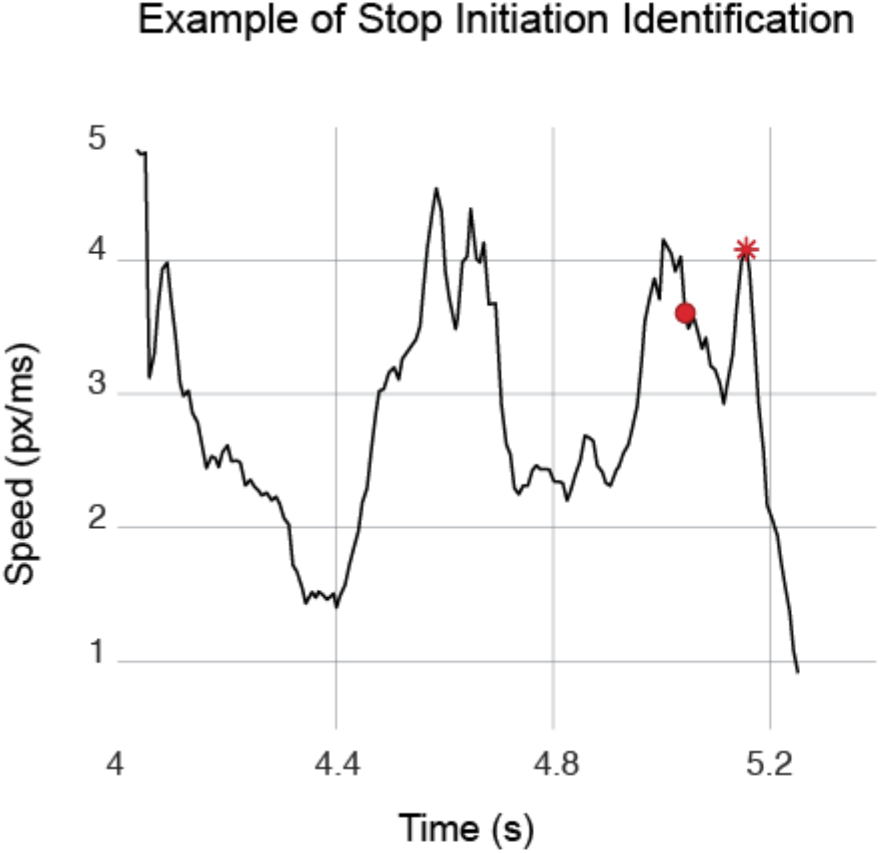
Example definition of stop initiation time. Circle indicates stop signal presentation. Star indicates stop initiation time.

### Statistics

#### Reaction Times and SSRTs

For each task version, average SSRTs for planned and unplanned stop trials were compared to one another using either a paired t-test for significance if distributions were normal or a Wilcoxon signed-rank if they were not, as determined using the Shapiro-Wilk test of normality. Additionally, mean SSRTs and RTs for participants from all four versions (*N* = 49) were combined and differences in these measures between trial types were evaluated using the same statistical tests. To identify any potential effect of task version on stopping behavior, a Bayesian ANOVA was performed. These tests were repeated to examine effects of condition and task version on RT.

#### Speed

We used a Wilcoxon signed-rank test to evaluate differences in percent change in speed and instantaneous speed at stop initiation between trial types because distributions of percent change in speed and instantaneous speed data did not meet assumptions of normality as determined using the Shapiro-Wilk test of normality. A paired samples t-test was used to evaluate differences in average speed across the entire trial, stop initiation time, and the difference in the time between stop initiation and SSRT for planned and unplanned stop trials. We used a Pearson correlation to assess the relationship between SSRT and (1) average speed across the entire trial, (2) stop initiation, and (3) instantaneous speed at stop initiation. A Spearman correlation was used to assess the relationship between SSRT and (4) percent change in speed.

## RESULTS

### Reaction Times

SSRTs were evaluated collectively (fig 6), combining all four versions of the task, and individually for each of the four different versions of the CMST (fig 7). Behavioral results are listed in **Table 2**. Results were similar across all versions. Considered collectively, SSRTs were significantly shorter for planned stop trials (*M* = 314.23, *SD* = 77.7) compared to unplanned stop trials (*M* = 495.34, *SD* = 103.37), *W* = 0, *p* < .001. RTs between conditions did not differ significantly (*M* _*planned*_= 291.6ms, *SD* = 137.67; *M* _*unplanned*_= 285.6, *SD* = 161.64), *W* = 740, *p* = .208. To determine the extent to which these data share a common distribution, we performed a Bayesian test of equivalence for RTs between the unplanned and planned conditions. With this approach the result was recapitulated, *BF*_10_ = .292. The difference in SSRTs for planned compared to unplanned stop trials was also consistent for all task versions when considered individually (as was the lack of a difference for the RT effects, **Table 2**, fig 7). Additionally, a Bayesian ANOVA revealed that there was no interaction effect between task version and trial type on SSRT or RT, *BF*_10 SSRT_ = .082, *BF*_10 RT_ = .002.

**Table 2.**
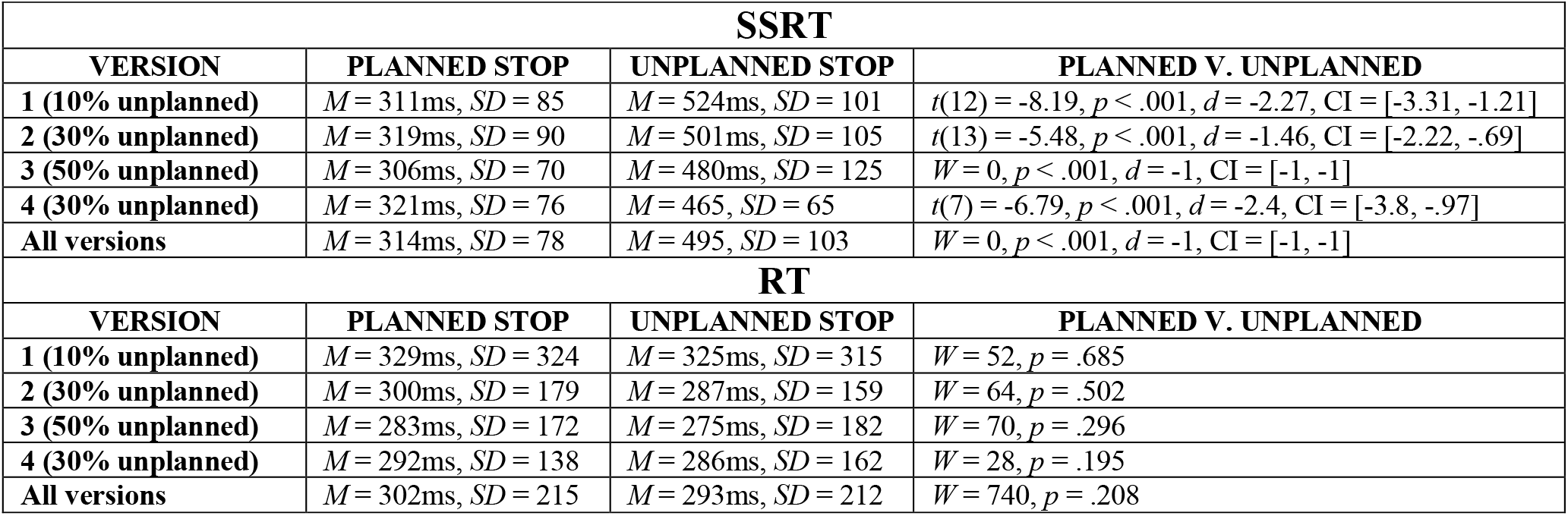
Behavioral results for task versions 1-4 and for all four versions combined. M = mean, SD = standard deviation. Effect size and 95% confidence intervals reported for all significant results.

| SSRT |  |  |  |
| --- | --- | --- | --- |
| VERSION | PLANNED STOP | UNPLANNED STOP | PLANNED V. UNPLANNED |
| 1 (10% unplanned) | $M = 311\text{ms}$, $SD = 85$ | $M = 524\text{ms}$, $SD = 101$ | $t(12) = -8.19$, $p < .001$, $d = -2.27$, $CI = [-3.31, -1.21]$ |
| 2 (30% unplanned) | $M = 319\text{ms}$, $SD = 90$ | $M = 501\text{ms}$, $SD = 105$ | $t(13) = -5.48$, $p < .001$, $d = -1.46$, $CI = [-2.22, -.69]$ |
| 3 (50% unplanned) | $M = 306\text{ms}$, $SD = 70$ | $M = 480\text{ms}$, $SD = 125$ | $W = 0$, $p < .001$, $d = -1$, $CI = [-1, -1]$ |
| 4 (30% unplanned) | $M = 321\text{ms}$, $SD = 76$ | $M = 465$, $SD = 65$ | $t(7) = -6.79$, $p < .001$, $d = -2.4$, $CI = [-3.8, -.97]$ |
| All versions | $M = 314\text{ms}$, $SD = 78$ | $M = 495$, $SD = 103$ | $W = 0$, $p < .001$, $d = -1$, $CI = [-1, -1]$ |
| RT |  |  |  |
| VERSION | PLANNED STOP | UNPLANNED STOP | PLANNED V. UNPLANNED |
| 1 (10% unplanned) | $M = 329\text{ms}$, $SD = 324$ | $M = 325\text{ms}$, $SD = 315$ | $W = 52$, $p = .685$ |
| 2 (30% unplanned) | $M = 300\text{ms}$, $SD = 179$ | $M = 287\text{ms}$, $SD = 159$ | $W = 64$, $p = .502$ |
| 3 (50% unplanned) | $M = 283\text{ms}$, $SD = 172$ | $M = 275\text{ms}$, $SD = 182$ | $W = 70$, $p = .296$ |
| 4 (30% unplanned) | $M = 292\text{ms}$, $SD = 138$ | $M = 286\text{ms}$, $SD = 162$ | $W = 28$, $p = .195$ |
| All versions | $M = 302\text{ms}$, $SD = 215$ | $M = 293\text{ms}$, $SD = 212$ | $W = 740$, $p = .208$ |

**Figure 6.**
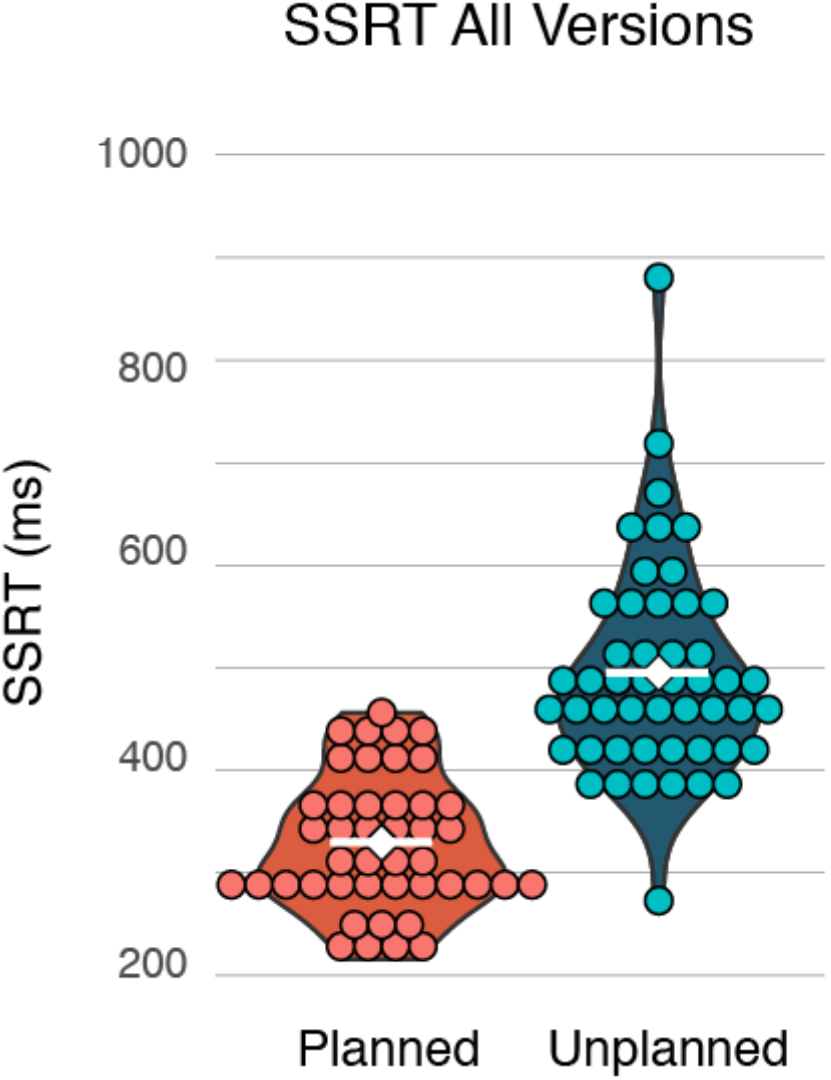
Combined SSRTs from all 4 task versions (N=49). Dots represent individual subjects. White diamond and line indicate group mean. SSRTs for planned stop trials (M = 314ms, SD = 78) were significantly shorter than unplanned stop trials (M = 495ms, SD = 103.37), W = 0, p < .001

**Figure 7.**
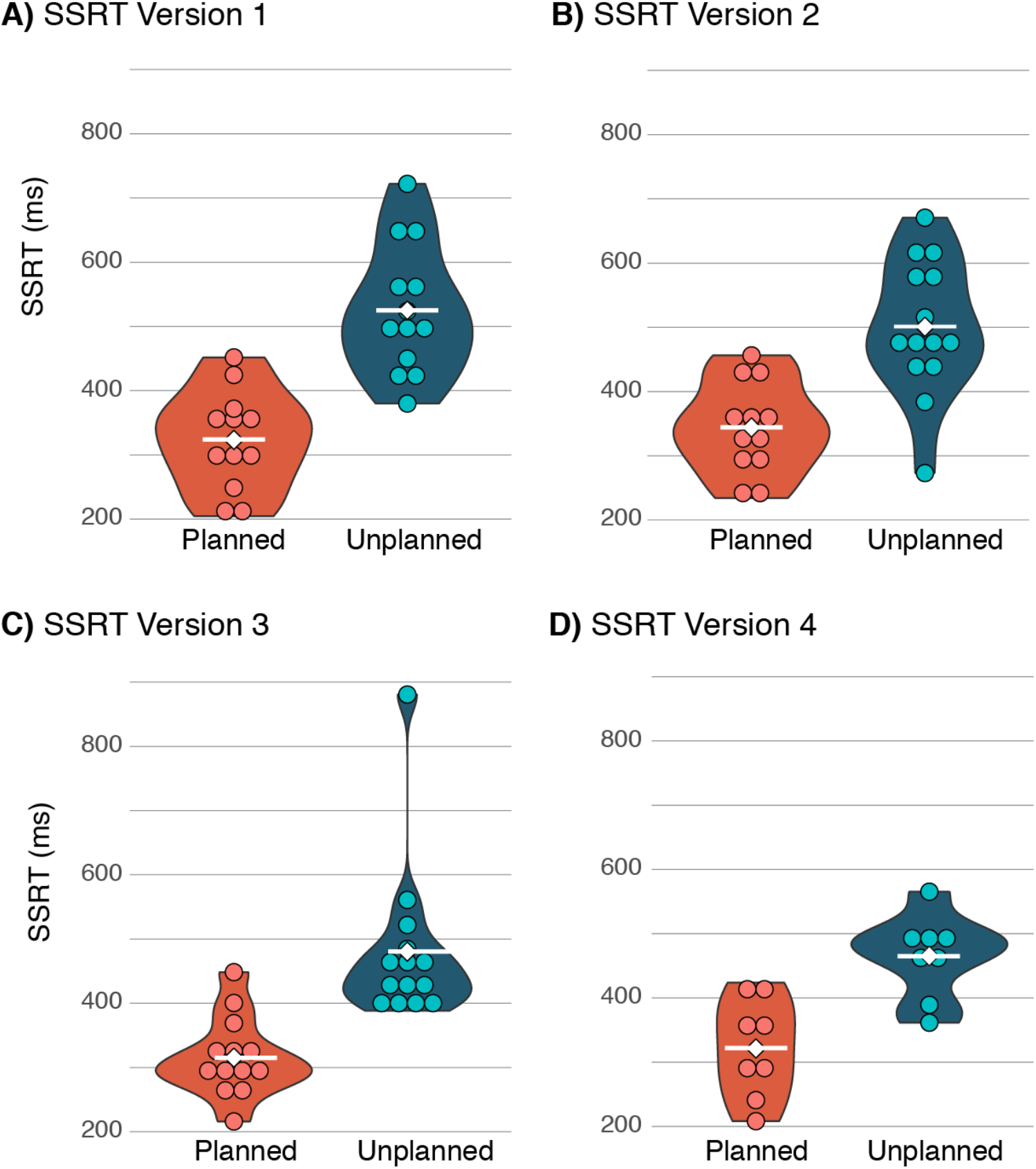
SSRTs for each task version. Dots represent individual subjects. White diamond and line indicate group mean. **A)** Version 1: 10% unplanned stop (N=13). SSRTs were significantly shorter for planned (M = 312ms, SD = 84.88) than unplanned stop trials (M = 525ms, SD = 101.04), t(12) = -8.19, p < .001. **B)** Version 2: 30% unplanned stop (N=14). SSRTs were significantly shorter for planned (M = 319ms, SD = 90.24) than for unplanned stop trials (M = 501ms, SD = 105.45), t(13) = -5.48, p < .001. **C)** Version 3: 50% unplanned stop (N = 14). SSRTs for planned stop trials (M = 306ms, SD = 69.56) were significantly shorter than those for unplanned stop trials (M = 480ms, SD = 125.02), W = 0, p < .001. **D)** Version 4: 30% unplanned with constraint (N = 8). SSRTs were significantly shorter for planned (M = 321ms, SD = 76.61) compared to unplanned stop trials (M = 465ms, SD = 64.62), t(7) = -6.79, p < .001.

### Speed

We found that participants slowed down significantly more on planned (*M* = -4.69, *SD* = 2.81) than on unplanned stop trials (*M* = 1.61, *SD* = 4 .14), *W* = 1, *p* = .016, *d* = -.944, CI = [-.988, -.762] (fig. 8a). Percent change in speed was significantly positively related to SSRT, *r* = .576, *p* = .022 (fig. 8b). That is, the more participants slowed their speed, the faster they stopped. Mean speed across the entire trial did not significantly differ between planned (*M* = 3.3px/ms, *SD* = .77) and unplanned stop conditions (*M* = 3.26 px/ms, *SD* = .8), *t*(7) = .946, *p* = .376. We did not find a correlation between trial average speed and SSRT, *r* = .096, *p* = .725.

**Figure 8.**
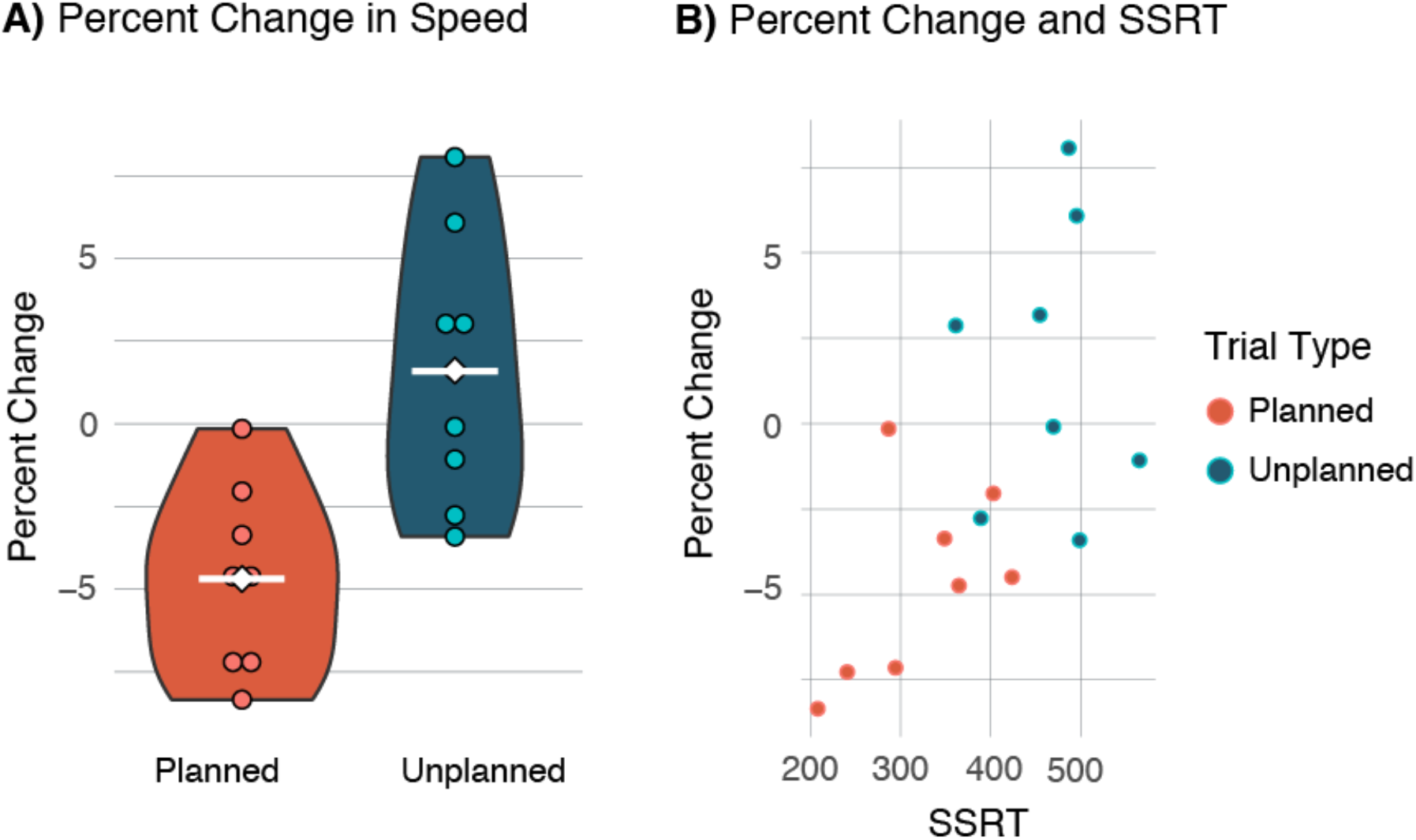
Percent change in speed between mid-trials movement and end of trial movement. **A)** Percent change in speed for each trial type. Dots represent individual subjects. White diamond and line indicate group mean. Wilcoxon signed-ranks test revealed that participants reduced speed significantly more on planned (M = - 4.69, SD = 2.81) than on unplanned-stop trials (M = 1.61, SD = 4.14), W = 1, p = .016. **B)** Percent change in speed was correlated with SSRT, r = .58, p = .02.

Stop initiation occurred significantly earlier on planned (*M* = 204ms after stop signal, *SD* = 75) compared to unplanned stop trials (*M* = 349ms after stop signal, *SD* = 77), *t*(7) = -6.451, *p* < .001, *d* = - 2.281, CI = [-3.621, -.905] (fig 9a). Stop initiation time was also significantly related to SSRT, *r* = 0.914, *p* < .001 (fig 9b). The time between stop initiation and SSRT was significantly longer for planned (*M* = 146.18ms, *SD* = 0.03) compared to unplanned stop trials (*M* = 126.63ms, *SD* = .03), *t*(7) = 3.56, *p* = .01, *d* = 1.258, CI = [.288, 2.183] (fig. 10). Instantaneous speed at the time of stop initiation was not significantly different between trial types (*M* _*planned*_ = 4.62px/ms, *SD* = 1.51; *M* _*unplanned*_ = 4.766px/ms, *SD* = 1.754), *t*(7) = -1.281, *p* = .241, and was not correlated with SSRT, *r* = .13, *p* = .641.

**Figure 9.**
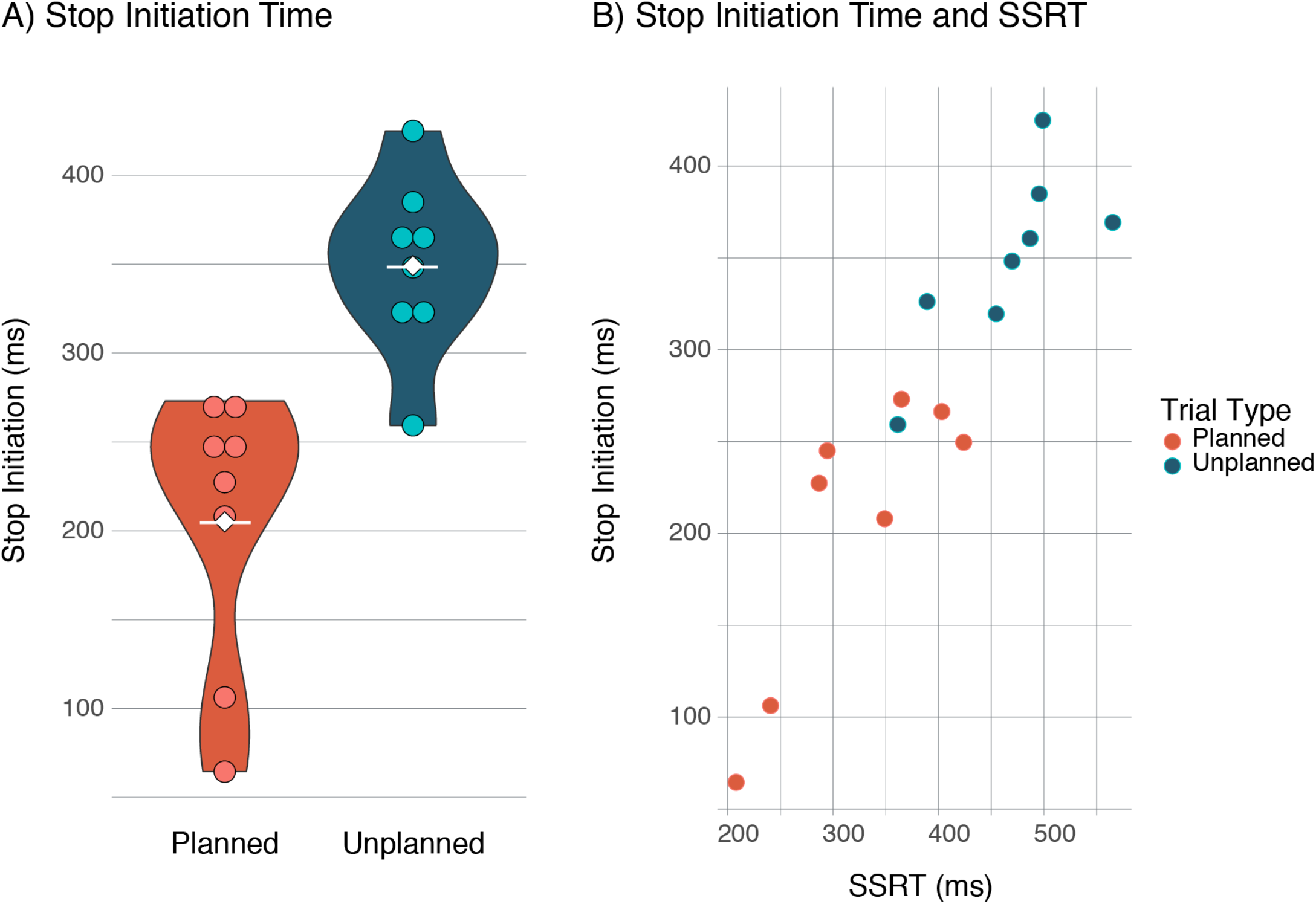
Stop initiation. **A)** Participants initiated stopping significantly earlier on planned (M = 205ms after stop signal, SD = .08) compared to unplanned stop trials (M = 349ms after stop signal, SD = .05), t(7) = -6.451, p< .001. Dots represent individual subjects. White line indicates group mean. **B)** Stop initiation time was significantly correlated with SSRT, r = .914, p < .001. Blue dots indicate planned stop trials. Orange dots indicate unplanned stop trials.

**Figure 10.**
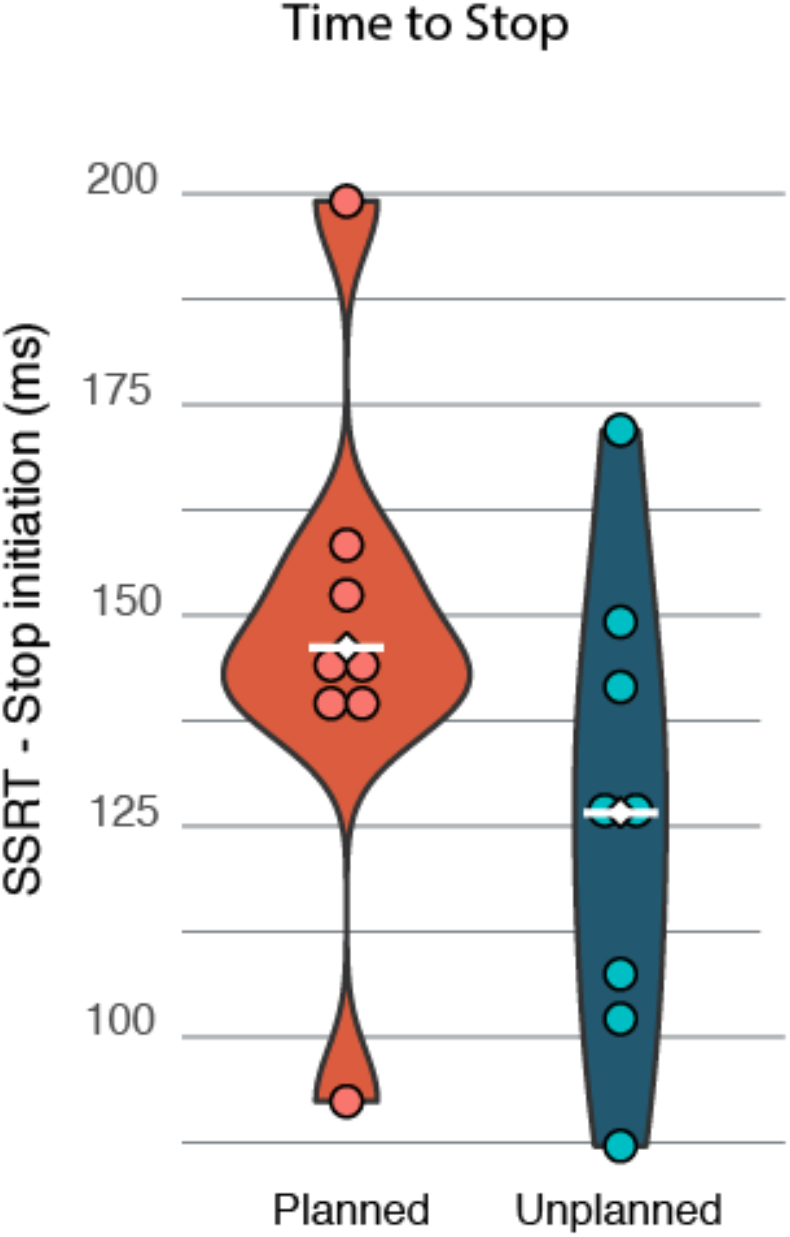
Time to stop. The difference between stop initiation time and SSRT was significantly greater for planned (M = 146ms, SD = .03) compared to unplanned stop trials (M = 127ms, SD = .03), t(7) = 3.558, p = .01.

## DISCUSSION

In the present study, we tested and evaluated a novel stop signal task that requires termination of a continuous movement. We found a consistent difference in SSRT between planned and unplanned stop trials across all versions of the task (fig 8), suggesting that the effect was robust, and that proportion of unplanned stop trials does not have a significant effect on the behavior. This is a beneficial feature of this task in that it allows for flexibility in total task length without compromising integrity or interpretability of the data. For example, a 50% unplanned stop version could be used in a shortened version of the task, reducing the temporal demand on the participants without compromising the ability to behaviorally differentiate planned and unplanned stopping. This could prove especially valuable for patient or pediatric populations.

Our finding that participants stopped earlier on planned compared to unplanned stop trials is consistent with existing literature concerning proactive inhibitory control and may be at least partially explained by proactive engagement of inhibitory mechanisms (Aron, 2011 for review). In previous research employing the SST, the degree of proactive inhibition (indexed by an increase in RT) was negatively correlated with SSRT, meaning that the more participants slowed their responses the faster they stopped (Aron et al., 2007; Jahfari et al., 2009). Although proactive inhibitory mechanisms are largely studied in the context of inhibiting a prepared discrete movement, it is possible similar mechanisms are involved in terminating an ongoing movement. Our observation that participants slowed prior to stop signal arrival significantly more on planned than unplanned stop trials also supports the possibility that participants engaged a breaking mechanism to facilitate faster SSRT on planned stop trials.

In addition to SSRT (defined by full motor arrest), we were able to capture behavioral changes reflecting initiation of the stop process. ‘Stop initiation’ was defined as the time point immediately preceding consistent deceleration eventuating in full motor arrest (fig. 5). We found that participants initiated stopping significantly earlier on planned than unplanned stop trials (fig. 9). Our data also revealed that the time between stop initiation and SSRT was longer for planned than unplanned stop trials (fig. 10). That is, even though participants stopped sooner on planned stop trials it actually took them longer to fully terminate their movement. This finding is consistent with what is predicted by the hypothesis that prepared inhibitory processes recruit the relatively slower indirect pathways while reactive inhibition relies more on the fast, mono-synaptic hyperdirect pathways (Aron, 2011 for review). Our observation that instantaneous speed at the time of stop initiation was not significantly different between trial types and was not related to SSRT suggests that a simple difference in speed cannot account for the discrepancy in the time it takes to stop between planned and unplanned stop trials.

There are other tasks that address some of the same limitations of the traditional SST that we sought to address here (Alegre et al., 2008; Hervault et al., 2019; Lofredi et al., 2020; Morein-Zamir et al., 2004, 2006; Morein-Zamir & Meiran, 2003; Slater-Hammel, 1960). Despite the availability of many versions of the stop signal task, the CMST is a valuable addition because it simultaneously allows for direct observation and precise comparison of the timing and mechanisms related termination of continuous movements under both prepared and reactive (i.e. planned versus unplanned stopping) conditions while providing the experimental control needed to be amenable to neuroscientific techniques and research in patient populations.

Use of the CMST during electrophysiological recordings and functional imaging will be a critical step toward confirming the generality of this putative neural inhibitory control system. If used in conjunction with neuroscientific techniques such as EEG or fMRI, this measure could provide a clearer picture of the timing and mechanisms underlying stopping than is currently available.

### Limitations

Although the present study represents an exciting next step in inhibitory control research, there are limitations that should be addressed in future research. One limitation is the small sample of participants who completed version 4, the only version for which we could confidently evaluate speed of movement. Although data collection was stopped prior to our target sample size of 12 (due to the COVID-19 pandemic), a post-hoc power analysis performed in G*power (Faul et al., 2007, 2009) revealed that our sample of *N* = 8 provided sufficient power, with α = 0.5, to detect the observed effect sizes for differences in SSRT (*power* = 0.99), stop initiation time (*power* = 0.99), and time to stop (*power* = 0.94). Power to detect the observed effect size for differences in percent change in speed between trial types, with α = 0.5, was 0.77, which is less than the recommended level of 0.8. Thus, it is possible that potentially informative differences in behavioral measures of speed changes were obscured by our sample size. Additionally, because larger movements could not be reliably captured with our set-up, we were not able to evaluate the possibility that magnitude of movement impacts stopping behavior. These limitations may be effectively addressed with behavioral constraints that are more compelling than a circular paper guide. One method we have found tentative success with is asking participants to use only their wrist when making the circular motions. Finally, our design is unable to disentangle the impact of temporal acuity on SSRT for planned stop trials. That is, variability in planned stop SSRT may be caused by an over- or under-estimation of the 1-second interval duration before the stop signal is presented rather than variability within the inhibitory control system itself.

### Conclusion

We found a reliable difference in SSRT for all task versions with subjects requiring less time to stop (shorter SSRTs) for planned compared to unplanned stop trials. Additionally, we found that there was more slowing and that stopping was initiated earlier on planned stop trials compared to unplanned stop trials. These data support the efficacy of the CMST for evaluating prepared and reactive termination of continuous movement (i.e. stopping that is planned vs unplanned). Finally, we found that it took participants longer to arrive at full motor arrest following stop initiation on planned compared to unplanned stop trials. Our CMST is an important addition to the field of inhibitory control research as it provides an unambiguous behavioral measure of stopping on individual trials, allows for direct comparison of proactive and reactive inhibitory processes, yields a rich set of behavioral measures to examine, and provides an opportunity to evaluate termination of a continuous movement-a currently understudied aspect of inhibitory control.

## ACKNOWLEDGEMENTS

We thank the Renée James Seed Grant to Initiative to Accelerate Scientific Research for support for this project. We also thank Drs. Ian Greenhouse and Jan Wessel for helpful comments on the manuscript.

